# Validation of a novel molecular host response assay to diagnose infection in hospitalized patients admitted to the ICU with acute respiratory failure

**DOI:** 10.1101/117853

**Authors:** Maria E. Koster-Brouwer, Diana M. Verboom, Brendon P. Scicluna, Kirsten van de Groep, Jos F. Frencken, Davy Janssen, Rob Schuurman, Marcus J. Schultz, Tom van der Poll, Marc J.M. Bonten, Olaf L. Cremer, on behalf of the MARS consortium

**Author notes:** **Corresponding author** Maria E. Koster-Brouwer, Julius Center for Health Sciences and Primary Care & Department of Intensive Care Medicine, University Medical Center Utrecht, Mailing address: Room F06.149, P.O. Box 85500, 3508 GA Utrecht, the Netherlands, T. E. The MARS Consortium also includes the following persons: Friso M. de Beer MD; Lieuwe D. J. Bos PhD; Gerie J. Glas MD; Roosmarijn T. M. van Hooijdonk MD, PhD; Janneke Horn MD, PhD; Mischa A. Huson MD; Peter Klein Klouwenberg MD, PhD; David Y. Ong MD, PhD; Laura R. A. Schouten MD; Marleen Straat MD; Lonneke A. van Vught MD, PhD; Luuk Wieske MD, PhD; Maryse A. Wiewel MD, PhD; Esther Witteveen MD. **Conflict of interest** The authors declare that they have no conflicts of interest related to the subject matter.

## Abstract

**Purpose:** The discrimination between infectious and non-infectious causes of acute respiratory failure (ARF) in hospitalized patients admitted to the intensive care unit (ICU) is difficult. Using a novel diagnostic test measuring the expression of four RNA biomarkers in blood (SeptiCyte LAB) we aimed to distinguish between infection and inflammation in this setting.

**Methods:** We enrolled hospitalized patients with ARF requiring prompt intubation in the ICU from 2011 to 2013. We excluded patients having an established infection diagnosis or an evidently non-infectious reason for intubation. Blood samples were collected upon ICU admission. Test results were categorized into four probability bands (with higher bands indicating a higher probability of infection) and compared with the plausibility of infection as rated by post-hoc assessment using predefined definitions.

**Results:** Of 467 included patients, 373 (80%) were treated for a suspected infection at admission. Plausibility of infection was classified as ruled-out, undetermined, or confirmed in 41 (11%), 135 (36%), and 197 (53%) of these, respectively. Overall, the pre-test probability of infection was 42%. Test results correlated with the plausibility of infection (Spearman’s rho 0.332; p<0.001). After exclusion of undetermined cases, positive predictive values were 29%, 54%, and 76% for probability bands 2, 3, and 4, respectively, whereas the negative predictive value for band 1 was 76%. However, SeptiCyte LAB did not outperform CRP when comparing diagnostic discrimination (AUC 0.731; 95%CI 0.677-0.786 vs. 0.727; 95%CI 0.666-0.788).

**Conclusion:** In a setting of hospitalized patients admitted to the ICU with ARF, the diagnostic value of SeptiCyte LAB seems limited.

## Introduction

Numerous biomarkers have been evaluated for diagnostic utility in distinguishing infection from sterile inflammation in critically ill patients, including C-reactive protein (CRP), procalcitonin (PCT), several coagulation markers, and others (1, 2). However, despite the clear association of these biomarkers with the presence of systemic inflammation, most did not diagnose or rule-out infection with sufficient rigor (1–4). Distinct protein biomarkers likely provide an (over)simplified representation of the host immune response to infection (2, 5), which is very complex yet largely similar to that following major surgery, trauma, and various other diseases triggering systemic inflammation (6). As a result, the use of single biomarkers may be predestined to yield only limited diagnostic value (2, 5).

Recently, a novel diagnostic test (SeptiCyte LAB, Immunexpress, Seatle, WA) was developed which aims to provide a probability of infection based on the expression of a specific genomic fingerprint consisting of CEACAM4 (carcinoembryonic antigen-related cell adhesion molecule 4), LAMP1 (lysosomal-associated membrane protein 1), PLA2G7 (phospholipase A2 group VII), and PLAC8 (placenta-specific 8-gene protein) (7). The simultaneous analysis of RNA transcription by these four genes in peripheral blood potentially utilizes information that is contained in various unrelated pathways of the host response at the transcriptome level. This new technology is recently approved by the American Food and Drug Administration (FDA), and is expected to be commercially available in the second half of 2017. In two technical validation studies SeptiCyte LAB was shown to be highly specific for infection in selected subgroups of both adult and pediatric patients – albeit including some subjects for whom the final diagnosis was already self-evident at the time of testing (7, 8). As a result, the precise clinical utility of the test for discriminating infectious and non-infectious causes of inflammation in the ICU remains unknown.

A population in which a diagnostic biomarker for infection would be particularly relevant includes patients admitted to the intensive care unit (ICU) with acute respiratory failure (ARF) after being in hospital wards. These patients are known to suffer from prolonged ICU stays and high mortality (9). Furthermore, dyspnea in these patients is a very non-specific symptom and its differential diagnosis is thus extensive, including congestive heart failure, pleural effusion, atelectasis, pulmonary embolus, acute respiratory distress syndrome, and—virtually always—infection. Early confirmation or rejection of infection is very relevant in this setting, since the former will secure timely initiation of antimicrobial therapy, whereas the latter might prompt a comprehensive diagnostic work-up for non-infectious (e.g. cardiac) causes of respiratory distress. Therefore, we aimed to determine the diagnostic and prognostic value of SeptiCyte LAB in patients admitted to the ICU from hospital wards with ARF.

## Methods

### Study design

For this nested cohort analysis, we selected patients who were enrolled in the Molecular Diagnosis and Risk Stratification of Sepsis (MARS) project, a prospective observational cohort study in two tertiary mixed ICUs in the Netherlands which started in 2011 (10). Participating centers were the University Medical Center Utrecht and the Academic Medical Center Amsterdam. Ethical approval for the study was provided by the Medical Ethics Committees of both participating hospitals, and an opt-out procedure to obtain consent from eligible patients was in place (protocol number 10-056).

### Patients

Patients were included if they had been admitted to the ICU between January 2011 and December 2013 with ARF (as evidenced by a need for mechanical ventilation within 24 hours of presentation) following admission to a hospital ward, coronary care unit, or medium care unit of at least 48 hours. Furthermore, all patients had to have an early warning score >5 (a clinical screening tool based on 6 cardinal vital signs (11)) and/or presence of ≥2 systemic inflammatory response syndrome (SIRS) criteria at ICU admission. Patients were excluded if they had another pertinent need for intubation and the ARF was evidently not caused by an infection (including, but not limited to, chronic respiratory insufficiency, primary cardiac arrest, and airway obstruction) or if a diagnosis of infection was already established at the time of ICU admission (i.e., infections for which antimicrobial therapy had been started >2 days prior to ICU admission).

### Reference diagnosis

Clinical data were prospectively collected by trained observers. Infectious events were registered using predefined definitions upon each occasion that antimicrobial therapy was initiated (10, 12, 13). For each event the plausibility of infection was categorized as none, possible, probable, or definite, based on a review of all available clinical, microbiological, and radiological data, and daily discussions with the attending team. For use as a reference test in the present analysis, we subsequently reclassified all events into the following discharge categories: infection ruled-out (patients with a post-hoc likelihood rated none, or patients who had never been treated for infection), infection undetermined (patients with possible infection), or infection confirmed (patients with a post-hoc likelihood rated probable or definite).

### SeptiCyte LAB tests

Blood specimens were collected within 24 hours of ICU admission in all patients using 2.5 mL PAXgene Blood RNA Tubes (PreAnalytiX GmbH, Hombrechtikon, Switzerland). According to the manufacturer’s instructions, samples were kept for a period of 2 to 72 hours at room temperature, and subsequently stored at-20°C (for a maximum of 1 month) and finally stocked at-80°C until analysis. RNA was then isolated on a QIAcube workstation using a PAXgene blood miRNA kit (Qiagen, Venlo, the Netherlands). The concentration of total RNA per sample was subsequently assessed by Nanodrop spectrophotometry (Agilent, Amstelveen, the Netherlands) and had to be between 2 and 50 ng/uL for a sample to be eligible for further analysis.

SeptiCyte LAB tests were performed in 96-well microtiter amplification plates on an Applied Biosystems 7500 Fast Dx Real-Time PCR Instrument (Thermo Fisher Scientific, Carlsbad, CA). During each amplification run, 3 control samples were also included. PCR results were initially quantified using ABI Sequence Detection Software version 1.4, after which data were imported into SeptiCyte Analysis software for interpretation. The SeptiCyte LAB score was calculated from the threshold cycle numbers (Ct-values) measured per gene as follows: *SeptiCyte LAB score = (Ct_PLA2G7_ + Ct_CEACAM4_) – (Ct_PLAC8_ + Ct_LAMP1_).* The resulting score (ranging between 0 and 10) was finally classified into four probability bands reflecting an increasing sepsis likelihood according to the manufacturer’s specification; scores ≤3.1 represented band 1 and were categorized as ‘sepsis unlikely’, whereas scores 3.1 to 4, 4 to 6, and >6 represented bands 2, 3, and *4,* respectively, and were categorized as ‘sepsis likely’.

### Statistical analysis

We performed mainly descriptive analyses to determine the value of SeptiCyte LAB in diagnosing infection, as formal assessment of test characteristics was precluded due to the relatively large proportion of patients in whom a final diagnosis of infection could not be made with certainty. However, some diagnostic measures were calculated, albeit not taking into account cases with an undetermined reference diagnosis. Furthermore, we calculated the area under the receiver-operating curve (AUROC) to compare the performance of the SeptiCyte LAB score with C-reactive protein (CRP), a biomarker commonly used in clinical practice. As CRP was not measured directly upon ICU admission in 115 (25%) cases, missing values were replaced with estimates derived from multiple imputation (details can be found in Appendix I) (14, 15).

To assess the potential utility of the SeptiCyte LAB score for risk stratification of patients upon ICU admission, we studied the relation of test results with case fatality (after correction for disease severity). For this purpose, we constructed two prognostic models, using the APACHE IV score either alone or combined with the SeptiCyte LAB score to predict 30-day mortality. Since the study had a multicenter design, we used generalized linear mixed models with a binomial distribution and logit link, and added a random intercept to accommodate possible outcome differences between both participating hospitals. Comparison of both models was based on Akaike’s information criterion (AIC) and the AUROC.

Differences between subgroups of patients were tested using the Wilcoxon Rank-Sum test or the Chi-square test, as appropriate. To test differences in patient characteristics associated with increasing SeptiCyte LAB scores, p-values for trend were calculated using the Cochran-Armitage trend test for dichotomous variables, or one-way ANOVA for continuous variables. Only if ANOVA suggested a significant association, then linear regression with the SeptiCyte probability band as group determinant was subsequently performed. All analyses were performed in SAS Enterprise Guide 4.3 (SAS Institute, Cary, NC) and R Studio (R Studio Team 2015, Boston, MA).

## Results

### Patients

Among 1399 hospitalized patients who were admitted to the ICU with ARF during the study period, 638 (46%) subjects were eligible for inclusion (Figure 1). Blood samples were unavailable in 157 of these, mostly because these specimens had been used for other studies within the MARS project. Additionally, 14 other patients had to be excluded due to technical issues during sample preparation or processing, leaving 467 (73%) subjects for final analysis. Compared to study patients, eligible patients without samples for analysis had less frequently congestive heart failure, more often chronic cardiovascular insufficiency, and higher APACHE IV scores and ICU mortality (Appendix II, eTable 1).

**Figure 1.**
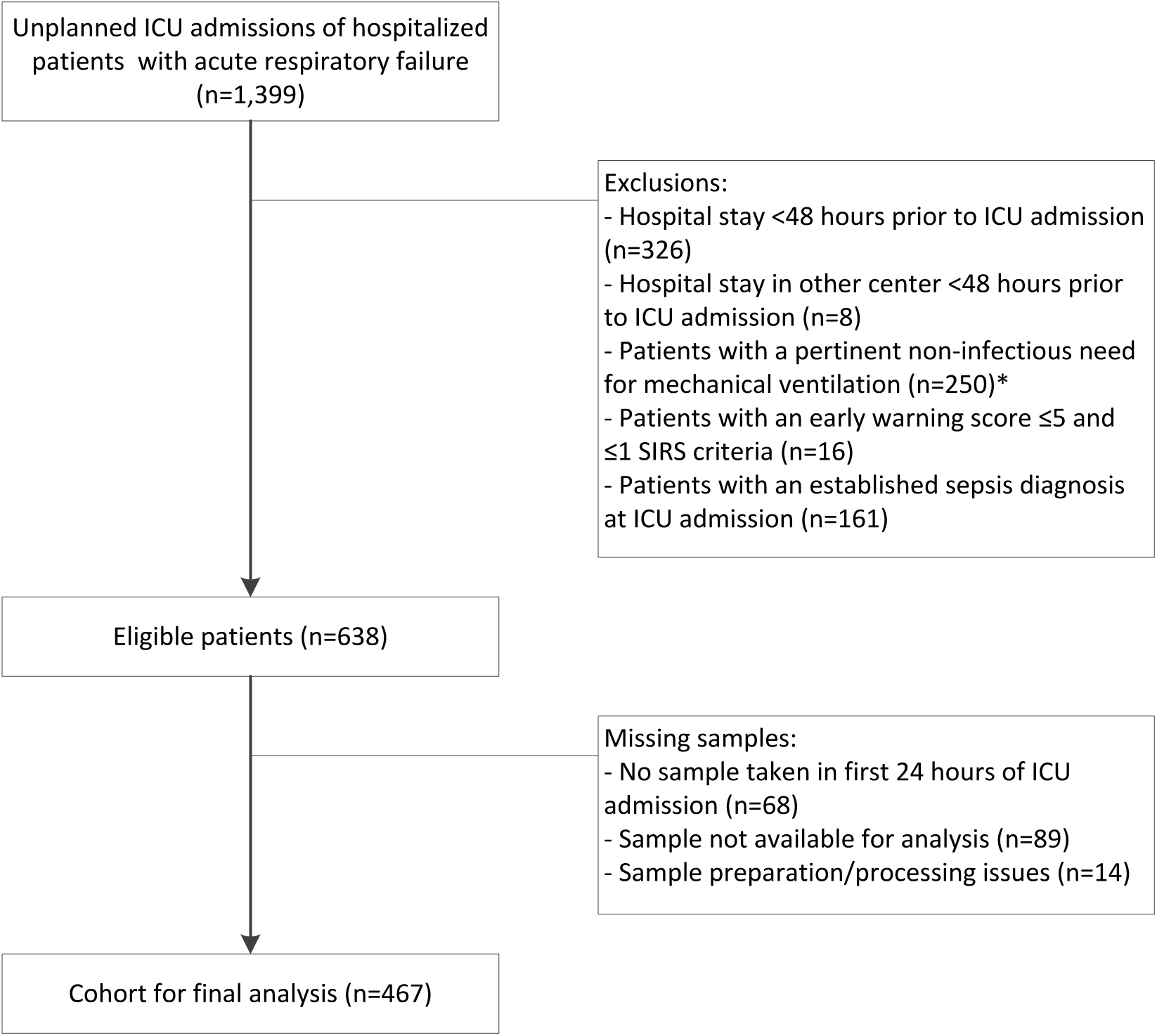
Flowchart of patient inclusions. ICU: intensive care unit. SIRS: systemic inflammatory response syndrome. * Including, but not limited to, patients with chronic respiratory insufficiency (n=107), in-hospital cardiac arrest (n=103), and airway obstruction (n=18).

**Table 1.**
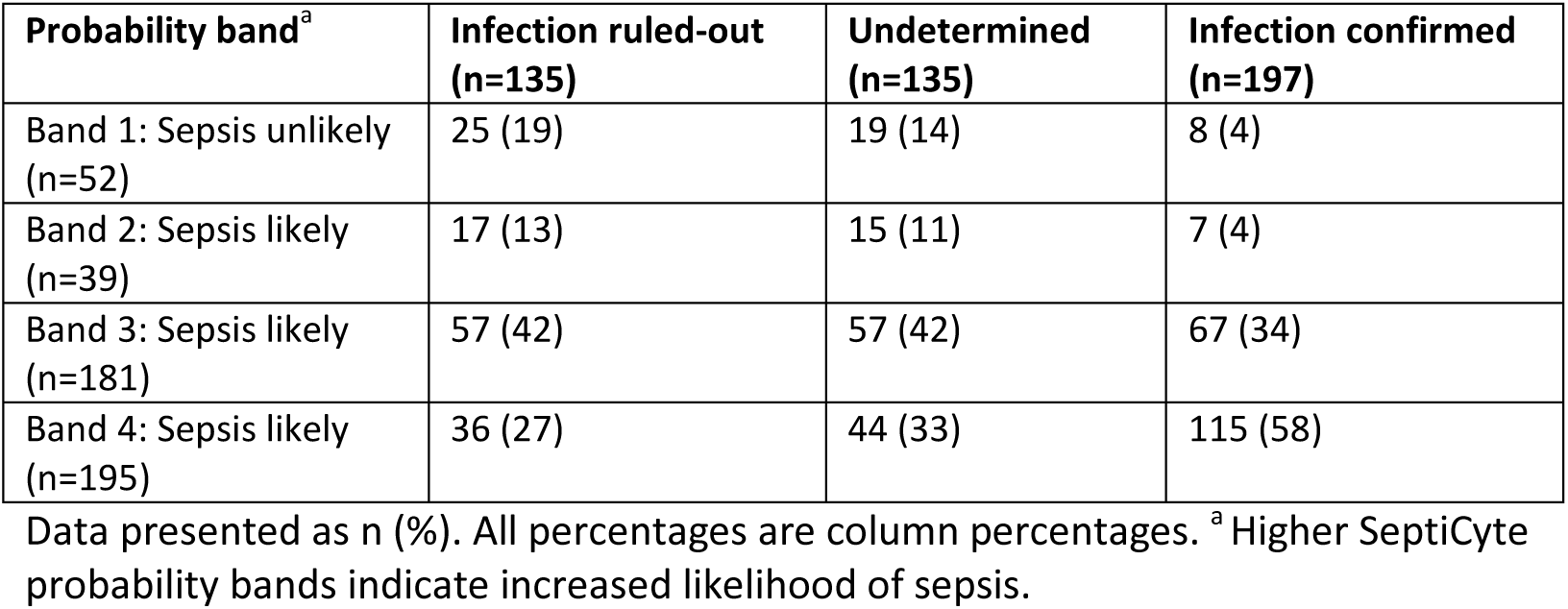
SeptiCyte LAB result versus reference diagnosis. Data presented as n (%). All percentages are column percentages. ^a^ Higher SeptiCyte probability bands indicate increased likelihood of sepsis.

| **Probability band<sup>a</sup>** | **Infection ruled-out**<br>**(n=135)** | **Undetermined**<br>**(n=135)** | **Infection confirmed**<br>**(n=197)** |
| --- | --- | --- | --- |
| Band 1: Sepsis unlikely<br>(n=52) | 25 (19) | 19 (14) | 8 (4) |
| Band 2: Sepsis likely<br>(n=39) | 17 (13) | 15 (11) | 7 (4) |
| Band 3: Sepsis likely<br>(n=181) | 57 (42) | 57 (42) | 67 (34) |
| Band 4: Sepsis likely<br>(n=195) | 36 (27) | 44 (33) | 115 (58) |

### Presence of infection

Because of presumed infection, 359 (77%) of the 467 included patients were treated with antimicrobial medications on day 1 in the ICU, and another 14 (3%) subjects started treatment on day 2. Among these, the post-hoc plausibility of infection was rated none in 41 (11%) cases. An additional 94 patients never received antimicrobial therapy, yielding a total of 135 subjects in the study cohort in whom infection was considered ruled-out. The remaining 332 patients were classified as undetermined (n=135) or infection confirmed (n=197). Hence, in the total study population, the pre-test probability of infection was 197/467 (42%). Of the patients in whom infection was undetermined or confirmed, the most commonly suspected sites of infection were respiratory tract infections (n=228), abdominal infections (n=52), and bloodstream infections (n=36).

### SeptiCyte LAB results

In patients in whom infection was ruled-out, undetermined, or confirmed the median (IQR) SeptiCyte LAB scores were 4.8 (3.7–6.1), 5.3 (3.9–6.4) and 6.5 (5.2–8.1), respectively (Figure 2). Formal analysis yielded a significant correlation between test scores and the probability of infection (Spearman’s rho 0.320; p<0.001).

**Figure 2.**
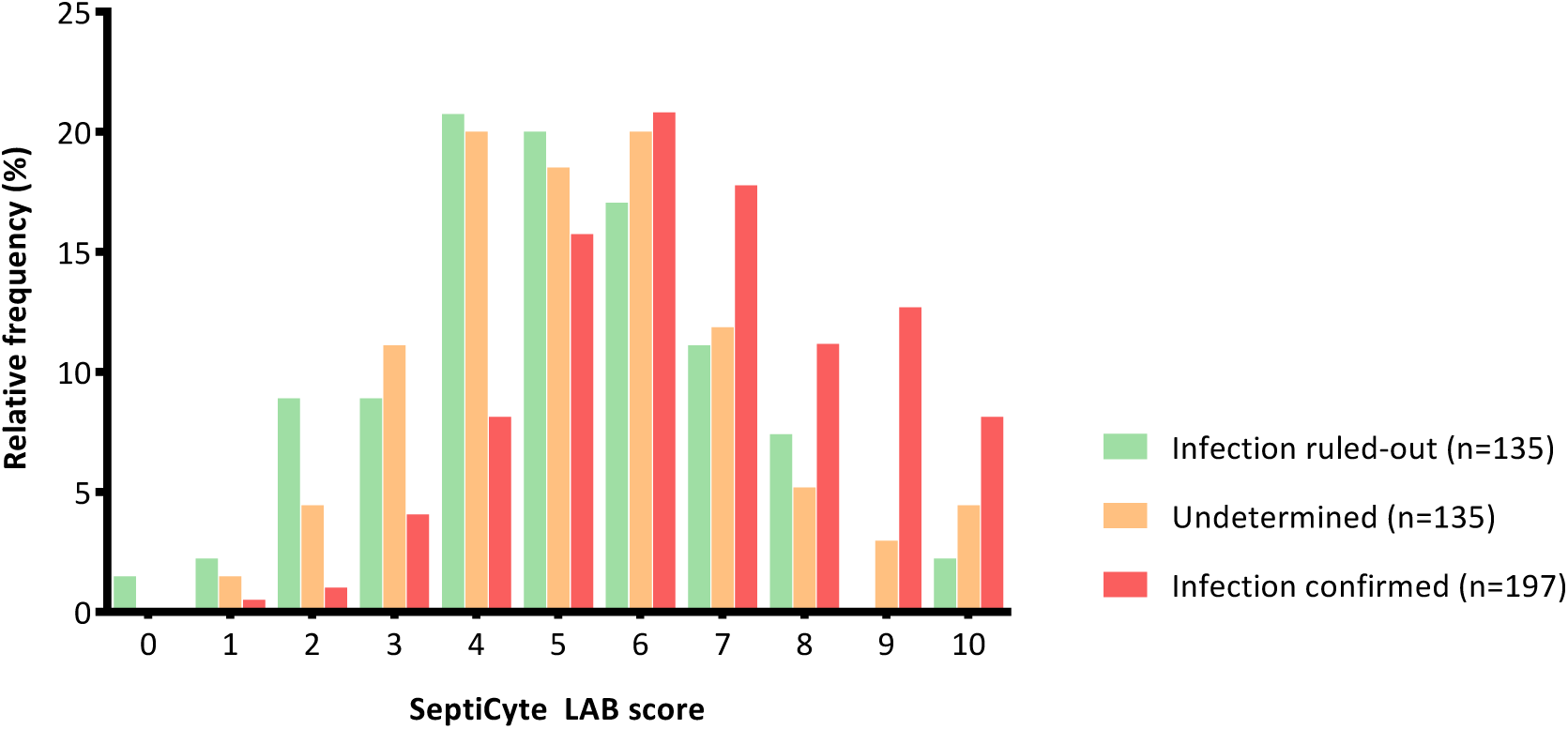
Distribution of SeptiCyte LAB scores by reference diagnosis. A higher SeptiCyte LAB score indicates a higher likelihood of sepsis. A score ≤3.1 should be interpreted as sepsis unlikely according to the manufacturer’s specification.

Table 1 shows the post-test probability bands for infection at admission according to the SeptiCyte LAB score in relation to the final (reference) diagnosis assigned during post-hoc assessment using all information available at ICU discharge. Concordance was observed in 189 (96%) of 197 patients with confirmed infection and in 25 (18%) of 135 patients in whom infection was ruled-out. After exclusion of undetermined cases, the positive predictive values for probability bands 2, 3, and 4 were 29%, 54%, and 76%, respectively. In contrast, the negative predictive value for probability band 1 was 76%.

### Discrepancy analysis

We observed 8 highly discordant cases where the test suggested that infection could be safely ruled-out, but in which infection was confirmed on post-hoc assessment. Review of these false negative patients revealed that they were older, had lower severity of illness upon presentation to the ICU, and more frequently had been previously admitted to the ICU than the 189 patients with a true positive test result (Table 2). Case descriptions for these patients are provided in Appendix III. Conversely, analysis of the 110 false positive patients showed that they had similar age, higher severity of illness, more previous ICU admissions, and were more likely to have been clinically suspected of infection than their 25 true negative counterparts. In addition, we compared Ct-values for individual RNA’s between false positive and true negative, and between false negative and true positive cases. Ct-values differed significantly for the PLA2G7, CEACAM4 and (possibly) PLAC8 genes, but not for LAMP1, when comparing false positive to true negative results (Figure 3-A). Similarly, PLAC8 and PLA2G7 seemed to offer some value in discriminating false negative from true positive test results, whereas LAMP1 and CEACAM4 did not (Figure 3-B).

**Figure 3.**
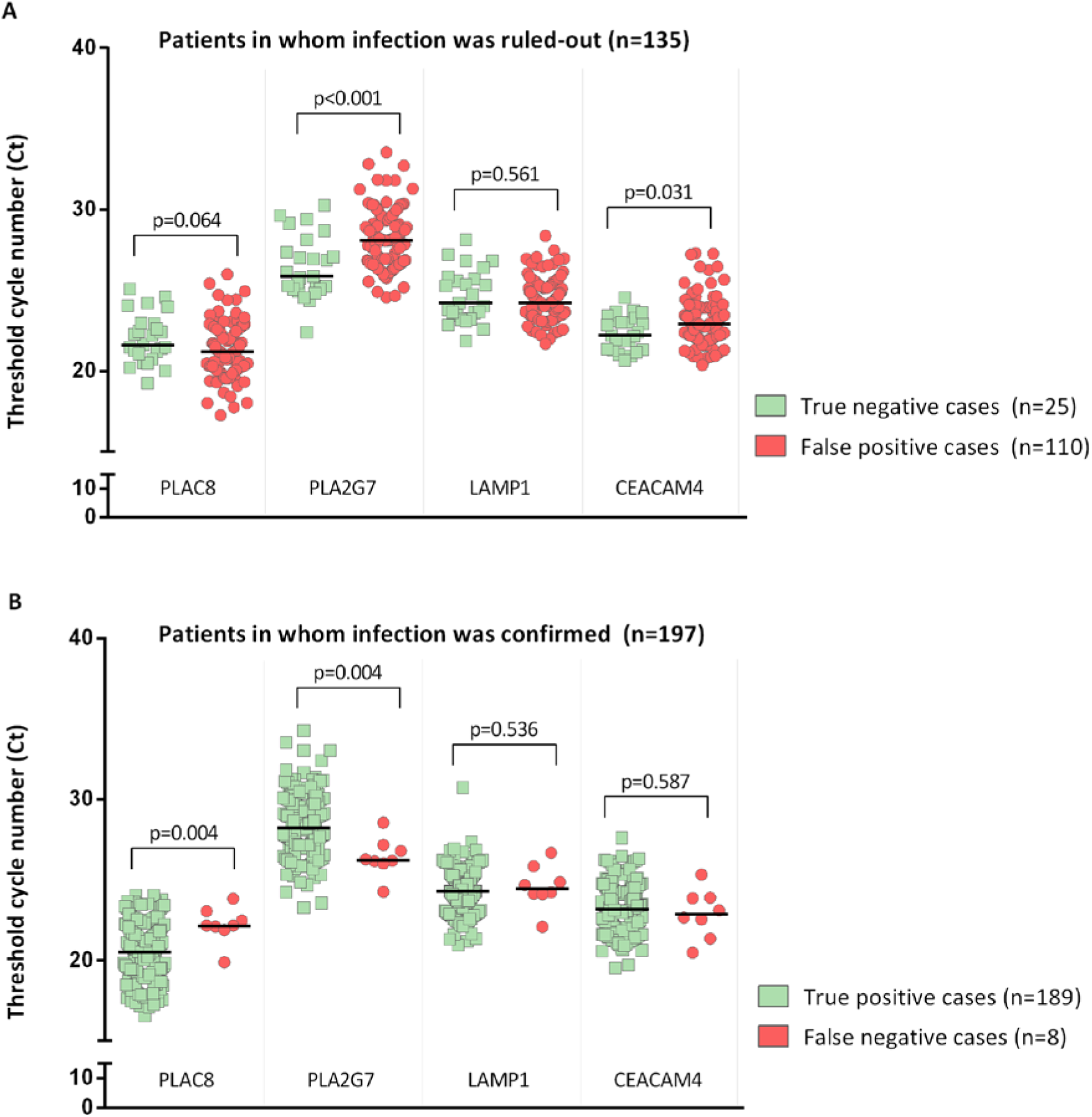
Median Ct-values per gene for non-infectious and infectious cases Accordinging to test result. For this analysis, cases in which infection was undetermined (n=135) were not taken into account as it was unknown whether the test classified them correctly.

**Table 2.** Characteristics of patients according to test result (discrepancy analysis) COPD: chronic obstructive pulmonary disease. CHF: congestive heart failure. APACHE: acute physiology and chronic health evaluation. ICU: intensive care unit. Continuous data are presented as medians (IQR), dichotomous data are presented as n (%). *Case descriptions for these patients are provided in Appendix III.

|  | **Infection ruled-out (n=135)** |  | **Infection confirmed (n=197)** |  |
| --- | --- | --- | --- | --- |
|  | **True negative** | **False positive** | **True positive** | **False negative\*** |
| Patients, N | 25 | 110 | 189 | 8 |
| Gender, female | 14 (56) | 48 (44) | 58 (31) | 1 (13) |
| Age, years | 60 (48-67) | 64 (53-75) | 65 (56-73) | 72 (64-77) |
| Surgical reason for admission | 1 (4) | 9 (8) | 9 (5) | 1 (13) |
| Comorbidities |  |  |  |  |
| - Congestive heart failure | 3 (12) | 10 (9) | 10 (5) | 0 (0) |
| - COPD | 3 (12) | 8 (7) | 23 (12) | 2 (25) |
| - Chronic cardiovascular insufficiency | 3 (12) | 1 (1) | 6 (3) | 1 (13) |
| - Diabetes mellitus | 4 (16) | 24 (22) | 36 (19) | 2 (25) |
| Immune deficiency | 3 (12) | 16 (15) | 30 (16) | 2 (25) |
| APACHE IV Score | 73 (60-86) | 81 (64-100) | 84 (70-101) | 64 (61-103) |
| Suspected site of infection |  |  |  |  |
| - Respiratory tract | 1 (4) | 20 (18) | 93 (49) | 6 (75) |
| - Abdominal | 0 (0) | 3 (3) | 40 (21) | 1 (13) |
| - Cardiovascular | 1 (4) | 3 (3) | 24 (13) | 1 (13) |
| - Other/unknown | 1 (4) | 12 (11) | 32 (17) | 0 (0) |
| - No suspicion | 22 (88) | 72 (65) | 0 (0) | 0 (0) |
| Prior ICU admission during hospital stay | 5 (20) | 57 (52) | 92 (49) | 6 (75) |
| ICU length of stay, days | 2 (1-8) | 4 (2-8) | 7 (3-12) | 4 (2-8) |
| ICU mortality | 2 (8) | 18 (16) | 36 (19) | 0 (0) |

### Comparative diagnostic evaluation

To better assess the clinical utility of SeptiCyte LAB, we compared its diagnostic performance to CRP. In patients in whom infection was later ruled-out, undetermined, or confirmed, median (IQR) plasma concentrations of CRP at ICU admission were 67 (22–152), 109 (63–207), and 166 (93–252) mmol/L, respectively. After exclusion of patients with an undetermined reference diagnosis, ROC analysis yielded an AUC of 0.727 (95%CI 0.666-0.788) for CRP versus 0.731 (95%CI 0.677-0.786) for SeptiCyte LAB (mean difference 0.004, 95%CI −0.077-0.086; p=0.919).

### Prognostic evaluation

Higher SeptiCyte LAB scores were associated with greater disease severity upon presentation to the ICU as well as with increased mortality (Appendix IV, eTable 2). However, a prognostic model that included both APACHE IV and SeptiCyte LAB scores was not superior in predicting 30-day mortality compared to a model using only the APACHE IV score (AUROC 0.737 versus 0.735, p=0.724; AIC 498 versus 497).

## Discussion

In this study, we evaluated the clinical utility of SeptiCyte LAB to diagnose infection in hospitalized patients that needed ICU admission because of ARF. Infectious episodes were correctly identified by the test in 96% of the patients with true infection. However, in patients in whom this diagnosis was refuted the test yielded a correct result in only 18% of patients. In fact, the test does not seem to offer better diagnostic discrimination in this setting than more commonly used biomarkers, such as CRP. In addition we assessed the prognostic value of SeptiCyte LAB, but when added to the APACHE IV model, the test did not improve prognostic power for predicting 30-day mortality.

Previous studies of SeptiCyte LAB have reported much higher discriminative power for infection than we found (AUCs of 0.88 and 0.99 vs. 0.73 in our study) (7, 8). The major difference between those reports and ours concerns the study domain. Early validation studies have mostly used cohorts in which infectious and non-infectious patients could clearly be distinguished on clinical grounds. For instance, one study compared children after cardio-pulmonary bypass surgery with severe sepsis cases (8). In another validation study of SeptiCyte LAB it was observed that the test performed better in a cohort of highly selected patients (AUC 0.95; 95%CI 0.91–1.00) than in a cohort representing more real-life settings (AUC 0.85; 95%CI 0.75-0.95) (7). Furthermore, a recent study investigating the diagnostic performance of SeptiCyte LAB across 39 publicly available datasets consisting of patients with either sterile inflammation or infection, reported highly variable findings (mean AUC 0.73, range 0.44-0.90 for individual datasets) (16). In search of a possible explanation for the lack of discriminative performance of SeptiCyte LAB in some cohorts, these authors noted that the expression of one of the four genes involved in calculation of the SeptiCyte LAB score (CEACAM4) was downregulated during sepsis only in the discovery cohort initially used to derive the score, but that no such evidence was observed in other cohorts (16). In our study, we found only minimal differences in gene expression of CEACAM4 between infectious and non-infectious cases, but this was also true for the other three genes (data not shown).

It is important to stress that in the current study we deliberately focused on a target population in which we knew beforehand that infection would be very difficult to diagnose with certainty. Many patients had significant (acute) comorbidities, had stayed in the hospital for prolonged periods of time prior to their ICU admission, and had previously been exposed to antibiotic treatment for (presumed) other infections. Furthermore, 171 of 638 eligible patients were excluded from analysis, mostly because their PAXgene samples had already been used for other studies within the MARS project. These patients frequently had a clinically apparent sepsis syndrome due to (confirmed) infection. These exclusions may thus have resulted in a study cohort that was more challenging to diagnose than a series of truly consecutive patients would have been. Nonetheless, one could argue there is little added value in using SeptiCyte LAB (or any other biomarker for that matter) in patients with clinically overt sepsis.

Although the probability of infection was prospectively adjudicated by trained observers, based on available post-hoc clinical, radiological and microbiological information, some diagnostic misclassification will most likely have occurred (10). This may have further reduced the apparent diagnostic performance of SeptiCyte LAB in our study. This stresses the difficulty of performing diagnostic studies in patients with infection, as a gold standard reference does not exist.

Nonetheless, the absence of a gold standard for infection cannot explain the observed differences with regard to discriminative power between our study and previous validation cohorts, nor the equipoise between SeptiCyte LAB and a more common host-response marker such as CRP in classifying infections (7).

In conclusion, in our clinical evaluation of SeptiCyte LAB in patients presenting to the ICU with ARF, who had already been hospitalized for other acute diseases and in whom infection thus was very difficult to diagnose, we could not replicate the high discriminative power that was formerly reported for this new biomarker. However, it is important to consider that SeptiCyte LAB scores are based on gene expression profiles which might vary between specific populations and/or settings (e.g., patients developing nosocomial infections following surgery, or patients with sepsis presenting to emergency departments). Therefore, more prospective research is needed before definite conclusions about the clinical utility of this novel test can be drawn.

## Acknowledgements

We thank Immunexpress for kindly providing SeptiCyte LAB kits and technical assistance.

## Electronic supplementary material

### Appendix I. Multiple imputation model and handling of imputed data

#### Multiple imputation model

As CRP was not measured at ICU admission in 115 (25%) cases, these missing values were replaced with estimates derived from multiple imputation. To impute these missing CRP values of the first whole day in the ICU we used an imputation method based on chained equations (package ‘mice’, version 2.25, 2015) (1). Variables that were used in the imputation model to predict the CRP value were: gender, age, race, medical or surgical admission, immune deficiency, Charlson comorbidity index, APACHE IV score, SOFA score at ICU admission, presence of infection at admission, SIRS criteria (fever, tachycardia, tachypnea, and abnormal white blood cell count) at ICU admission, ICU length of stay, and the SeptiCyte LAB score. If needed, rounding and boundaries were used to assure that clinically possible values were replaced for missing values. We performed 30 iterations and 25 imputations (3–5).

#### Handling imputed data

We averaged all AUROCs for CRP estimated from the imputed datasets in order to get an estimate for the discriminative power of CRP. Also, the difference in AUCs was calculated as the average difference between the AUROCs of SeptiCyte and CRP over all imputed datasets. Using Rubin’s rules, we calculated the accompanying 95% confidence intervals and test-statistics for these estimations to arrive at correct effect estimates and standard errors (6–7).

## Appendix II. Patient characteristics (eTable 1)

**eTable 1.** Characteristics of in- and excluded patients. COPD: chronic obstructive pulmonary disease. CHF: congestive heart failure. APACHE: acute physiology and chronic health evaluation. ICU: intensive care unit. Continuous data are presented as medians (IQR), dichotomous data are presented as n (%).

|  | **Exclusions**<br>**(patients without sample**<br>**and/or test result, n=171)** | **Inclusions**<br>**(patients with test result,**<br>**n=467)** | **P-value** |
| --- | --- | --- | --- |
| Gender, female | 66 (39) | 169 (36) | 0.577 |
| Age, years | 65 (54-73) | 65 (54-73) | 0.870 |
| Surgical reason for admission | 6 (4) | 25 (5) | 0.337 |
| Comorbidities |  |  |  |
| - Congestive heart failure | 4 (2) | 33 (7) | 0.024 |
| - COPD | 20 (12) | 52 (11) | 0.843 |
| - Chronic cardiovascular insufficiency | 13 (8) | 17 (4) | 0.036 |
| - Diabetes mellitus | 24 (14) | 88 (19) | 0.157 |
| Immune deficiency | 33 (19) | 68 (15) | 0.147 |
| APACHE IV Score | 87 (70-111) | 81 (65-98) | 0.010 |
| Admission diagnosis (top 3 categories) |  |  | 0.098 |
| - Pneumonia | 38 (22) | 131 (28) |  |
| - Congestive heart failure | 13 (8) | 34 (7) |  |
| - Atelectasis | 12 (7) | 32 (7) |  |
| Prior ICU admission during hospital stay | 75 (44) | 227 (49) | 0.287 |
| Intubation | 137 (80) | 394 (84) | 0.203 |
| ICU length of stay, days | 4 (2-11) | 5 (2-10) | 0.104 |
| ICU mortality | 42 (25) | 74 (16) | 0.012 |

## Appendix III. Case vignettes of false negative cases

### Patient #1

Patient had been previously admitted because of bilateral pneumonia. After 2 weeks she was readmitted to ICU with respiratory insufficiency mainly due to fluid overload and atelectasis. However, the patient was also suspected of recurrent pneumonia and because of earlier growth of Enterobacter cloacae, she was treated with flucloxacillin, vancomycin, and ciprofloxacin. Sputum cultures subsequently grew Staphylococcus aureus.

### Patient #2

Patient had been treated for mediastinitis following a spontaneous retropharyngeal abscess for 6 weeks; 5 days after discontinuation of antimicrobial therapy he was readmitted to the ICU with tachypnea and fever. CRP 31/93. The differential diagnosis on admission included HAP, pulmonary embolism, abscess in neck/mediastinum, and empyema. CT-scan showed multiple mediastinal fluid collections for which percutaneous drainage was subsequently performed and ceftriaxon plus metronidazol was empirically started.

### Patient #3

Patient had been admitted for 5 days following esophagectomy. After 3 months he was readmitted with dypnea, fever and back pain. CRP 79/193. PCT 2/4. The differential diagnosis on admission included HAP, pulmonary embolism, and late anastomotic leakage. Chest X-ray showed both infiltrates and spinal fractures. Treatment was started with ceftriaxon. Blood cultures subsequently grew Klebsiella pneumoniae.

### Patient #4

History of DM induced nephropathy, for which continuous ambulatory peritoneal dialysis. Patient had been treated for 3 months with intraperitoneal vancomycin for catheter peritonitis (due to corynebacterium) until 4 months before ICU admission and had undergone CABG 1 month before ICU admission. He was readmitted with shock, respiratory insufficiency and left-sided pleural effusion. CRP 234. The differential diagnosis on admission included pneumosepsis, abdominal sepsis (due to recurring CAPD peritonitis), bowel ischemia, hematothorax. A laparotomy was performed, but was negative. Sputum cultures grew Klebiealla pneumoniae and Serratia marcescens. The patient recovered following ciprofloxacin treatment.

### Patient #5

Patient had been admitted for 5 days following pancreatectomy. After 3 weeks he was readmitted with hypothermia, lactic acidosis and hypercapnia. CRP 55. The differential diagnosis on admission included anastomotic leakage, abdominal abscess, pleural effusion, and exacerbation COPD. Laparotomy showed small bowel perforation with an infected pocket in the right upper quadrant. Patient was treated with ciprofloxacin/clindamycin/anidulafungin.

### Patient #6

History of laryngectomy. Patient underwent uncomplicated mandibular resection. 3 days later he was admitted to the ICU with hypoxia and (some) fever (38.2). CRP 76/155. PCT 0/0. The diagnosis was evident: massive aspiration pneumonia due to malfunction of his pre-existing (one-way) tracheal-oesophageal speaking valve. He was treated empirically with ceftriaxon.

### Patient #7

Immunocompromised patient following allogenic stem cell transplant for CML. Medical history of recurrent infections, most currently cellulitis of the left lower leg for which he was still receiving flucloxacillin. Presentation to ICU with hematemesis (most likely due to GvHD) and a new infiltrate. Patient was treated empirically with ceftazidim, clindamycin and voriconazol. Neither BAL nor blood cultures (which were both performed while patient was receiving antimicrobial treatment) yielded a definite causative pathogen.

### Patient #8

Patient had been admitted for 4 days following acute subdural hematoma (while on anticoagulation) for which surgical decompression had been performed. 10 days later he was readmitted with fever, new-onset atrial fibrillation, renal insufficiency, and respiratory distress. CRP 112/185. The differential diagnosis on admission included aspiration pneumonia (due to difficulty swallowing), wound infection (he had a lesion on the back of his head), secondary peritonitis, and decompensated heart failure. Patient was empirically treated with ceftriaxon (for possible pneumonia) and flucloxacillin (for presumed wound infection), but later switched to ciprofloxacin when the sputum and blood cultures both grew Serratia marcescens.

## Appendix IV. Patient characteristics by SeptiCyte LAB result (eTable 2)

**eTable 2.** Patient characteristics by SeptiCyte LAB result. Continuous data are presented as medians (IQR), dichotomous data are presented as n (%).

| Patient characteristics | SeptiCyte probability band |  |  |  | P-value for trend |
| --- | --- | --- | --- | --- | --- |
|  | Band 1 (n=52) | Band 2 (n=39) | Band 3 (n=181) | Band 4 (n=195) |  |
| Gender, female | 21 (40) | 13 (33) | 67 (37) | 68 (35) | 0.562 |
| Age | 67 (54-76) | 65 (53-75) | 63 (53-73) | 65 (55-73) | 0.710 |
| Immune deficiency | 7 (13) | 5 (13) | 21 (12) | 35 (18) | 0.253 |
| APACHE IV score | 74 (63-96) | 69 (59-89) | 80 (64-96) | 84 (70-102) | <0.001 |
| Prior ICU admission during hospital stay | 17 (33) | 22 (56) | 97 (54) | 91 (47) | 0.352 |
| ICU length of stay, days | 5 (2-9) | 4 (2-8) | 6 (3-11) | 6 (3-10) | 0.306 |
| ICU mortality | 4 (8) | 6 (15) | 24 (13) | 40 (21) | 0.020 |
| 30-day mortality | 7 (14) | 11 (28) | 37 (20) | 46 (24) | 0.271 |

